# Oligonucleotide Library Assisted Sequence Mining Reveals Promoter Sequences With Distinct Temporal Expression Dynamics For Applications In *Curvibacter* sp. AEP1-3

**DOI:** 10.1101/2024.03.24.586450

**Authors:** Maurice Mager, Lukas Becker, Nina Schulten, Sebastian Fraune, Ilka M. Axmann

**Author notes:** contributed equally (Maurice Mager, Lukas Becker).

## Abstract

The *ß-proteobacterial* species *Curvibacter* sp. AEP1-3 is a model organism for the study of symbiotic interactions as it is the most abundant bacterial colonizer of the basal metazoan *Hydra vulgaris*. Yet, genetic tools for *Curvibacter* are still in an infancy: few promoters have been characterized for *Curvibacter*. Here we employ an oligonucleotide based strategy to find potential expression systems derived from the genome of *Curvibacter*. Potential promoters were systematically mined from the genome in silico. The sequences were cloned as a mixed library into a mCherry reporter gene expression vector and single positive candidates were selected through Flow Cytometry based sorting to be further analyzed through bulk measurements. From 500 candidate sequences, 25 were identified as active promoters of varying expression strength levels. Bulk measurements revealed unique activity profiles for these sequences across growth phases. The expression levels of these promoters ranged over two orders of magnitudes and showed distinct temporal expression dynamics over the growth phases: while 3 sequences showed higher expression levels in the exponential phase than in the stationary phase, we found 12 sequences saturating expression during stationary phase and 10 that showed little discrimination between growth phases. From our library, promoters the genes encoding for DnaK, RpsL and an AHL synthase stood out as the most interesting candidates as their expression profiles fit a variety of applications. Examining the expression levels of successful candidates in relation to RNAseq read counts revealed only weak correlation between the two datasets. This underscores the importance of employing comprehensive high-throughput strategies when establishing expression systems for newly introduced model organisms.

## 1 Introduction

*Curvibacter* sp. AEP1-3 (hereafter *Curvibacter*) is a rod-shaped *ß-proteobacterial* species best known for its symbiotic interaction with *Hydra vulgaris* [1], a freshwater polyp of the basal metazoan family *Cnidaria*, a sister group to the *Bilateria*. Together with other members of *Hydras* microbiota, they form a complex system of bacteria-bacteria as well as bacteria-host interactions [2, 3, 4]. While the host provides an ecological niche to its colonizers, the microbiota affects mobility, asexual reproduction as well as protection against fungi [1]. This symbiotic relationship provides an invaluable avenue for the study of inter kingdom interactions and allows for the exploration of general principles of symbiosis in the natural world such as the remarkable host-microbe communication between *Curvibacter* and *Hydra vulgaris* established through the exchange of N-acetyl homoserine lactone [2].

The limited genetic accessibility of *Curvibacter* restricts advancements in genetically manipulating its cells, thereby impeding progress in the field of interkingdom symbiosis. Genomic modifications over homologous recombination [5] are cumbersome [6] and have a low success rate. *Curvibacter* cells are amenable to transformation using RSF1010 vector [7] constructs through conjugation with *E. coli* donor cells but only a limited number of promoters are accessible for use and none of them have been characterized to date. In this study, we set out to develop a strategy to create tool kit of viable promoters for *Curvibacter* to promote its use as a model organism.

The Anderson Collection of Synthetic Promoters is a good reference for the development of orthogonal constitutive expression systems (http://parts.igem.org/Promoters/Catalog/Anderson). The collection provides a range of expression levels that have been well characterized in many species such as *E*.*coli, V. natriegens* and some *Cyanobacteria* [8, 9] but as orthogonal promoters they usually show the same temporal expression dynamics in the form of a stable, constant activity over growth conditions, providing the same transcript level over the entire time of cultivation. However, when expressed from a plasmid, the total expression level of most promoters often increases significantly in the stationary phase due to changes in the copy number of most plasmids: the copy number generally increases with slower growth during stationary phase, leading to higher transcript expression and increased protein levels [10, 11, 12]. While this may be either desirable for some applications or irrelevant in experimental setups where cells are only observed during logarithmic growth, such accumulation may prove detrimental, for example, in long-term experiments. Therefore, for the development of plasmid-based expression systems with stable expression in different growth phases, orthogonal promoters that cannot absorb this burst of expression may not be the most suitable.

While hand picking or designing individual sequences with a predicted expression strength is a valid strategy to develop expression systems, the use of entire sequence libraries provides a promising alternative due to the high throughput of tested sequences [13]. Such libraries are generated by synthesizing oligonucleotide sequences on highly sophisticated commercial DNA synthesis platforms that allow the simultaneous generation of many sequences at the same time. These libraries are collected and purified in a single sample and can be used for cloning applications to create a library of plasmids each containing a different synthesized sequence, as well as, in our case, a reporter sequence such as the Green Fluorescence Protein (GFP) or mCherry. Downstream, the use of flow cytometry and cell sorting can aid in picking positive candidates from such libraries to avoid the extensive effort of manually picking and analyzing individual colonies. The aforementioned libraries can consist of sequences with varying degrees of randomization, generated by using mixed nucleotides during synthesis which allows for the incorporation of any base by chance [14]. This strategy is necessary for projects in which the investigators aim to obtain the best suited sequences from a bias free sequence space or if there is no information available that could reduce the degree of freedom. The GeneEE library of Lale *et al*. [15] follows exactly this approach by using long stretches of randomized nucleotides to find novel promoter sequences de novo.

An alternative we employ here is the generation of highly curated libraries. Limiting the pool only to sequences with a high probability of success simplifies downstream processes and can yield many more positive candidates in significantly smaller libraries, making it an easier and more cost efficient method. The generation of such libraries can be facilitated by neural networks, trained on existing promoter sequences, extracted from curated libraries such as the Prokaryotic Promoter Database (PDD) [16] or as in our case simply by using existing sequences harvested directly from the target species genome [17]. The latter approach results in finding promoter sequences that won’t be orthogonal, but it is a valid approach to also find expression systems which are either inducible or show a desired temporal expression dynamic in certain growth phases. Moreover, these sequences already inherit the genomic context for specific regulation, to a degree. By extracting those sequences, it allows the identification of different regulatory elements depending on the extracted length and culture conditions.

Aim of this study is the discovery and characterization of novel expression systems for the use in expression vectors for *Curvibacter*, with a special focus on promoters that provide stable expression independently on growth phases in liquid media. To increase the odds of individual promoter sequences, promoter and ribosomal sequences were not further discriminated, but entire 5^*′*^ upstream non-coding regions were used instead. Candidate sequences for this study were directly harvested (Figure 1) from the *Curvibacter* genome sequence (GCF_002163715.1). Sequences were ordered as single stranded oligonucleotides and cloned into an expression vector with mCherry as reporter gene. mCherry expression serves here as quantifiable proxy for expression strength of a given candidate sequence. Positive candidates were selected with the aid of flow cytometry and cell sorting and were subsequently analyzed via bulk fluorescence measurement. The extracted 5^*′*^ upstream non-coding regions displayed a wide range of activity levels that can be used for different applications. We discovered different behavior of the selected candidates in terms of their activity over growth phases: while most candidate promoters showed typical increased activity in late exponential- to stationary phase, we also found some promoters to be slightly more active in exponential than in stationary phase. Several sequences were found to show similar expression levels during exponential-as well as stationary phase. We mapped the strength of our characterized expression systems against normalized RNAseq read counts and found weak correlation between the two data sets, showing that RNAseq does not generally serve as a reliable tool to predict promoter strength, at least in an *in trans* context, showing how useful oligonucleotide library based approaches are to find promoter sequences with desired features.

**Figure 1:**
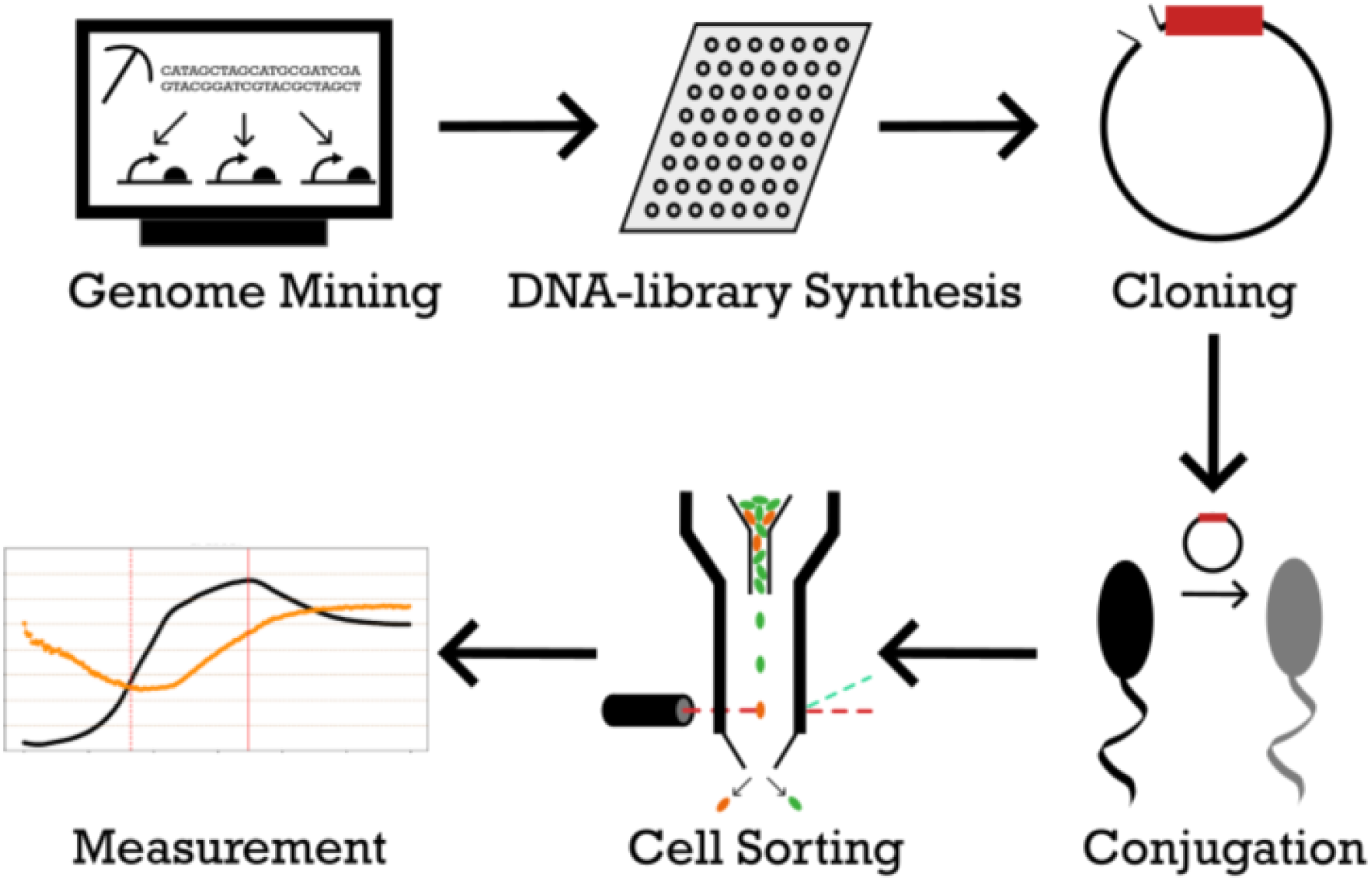
Schematic overview of the applied workflow. Design of the oligonucleotide library starts by mining the host genome for suitable promoter sequences. Sequences are synthesized, cloned and imported into the host species and sorted for activity by Flow Cytometry Cell Sorting. In depth assays of individual sequences characterize each candidate promoter in detail.

## 2 Material & Methods

### 2.1 Extraction of Promoter Sequences in the Genome of *Curvibacter* sp. AEP1-3

To extract candidate promoter sequences suitable for our experiments, first we retrieved the GenBank file for the *Curvibacter* genome from the GenBank FTP site, serving as our primary source of genomic data. All subsequent data processing and analysis was performed in the Python 3 programming environment within the Jupyter Notebook development platform. To extract and manage genomic data, we utilized the BioPython library for parsing the GenBank file. The scripts and all relevant data files can be found on our github repository (https://github.com/Kanomble/curvibacter_promotor_studies).

Local start and stop positions as well as the genomic orientation of all sequence elements labeled as “genes” within the *Curvibacter* genome were structured into a dictionary. Gene sequence identifiers were employed as keys but simultaneously also stored in a list object, preserving their exact positions within the genome. This list was instrumental for tracking sequence location and order. To provide context for the gene sequences, we implemented a parsing process to retrieve information about the current gene sequence, the previous gene sequence, and the next gene sequence based on the current identifier. To refine our dataset, we filtered out sequences that didn’t meet length criteria, specifically sequences with more than 170 base pairs and less than 60 base pairs, ensuring that only appropriately sized 5^*′*^ upstream non-coding regions were included in the analysis. Further filtering steps involve removing sequences with opposing orientations and overlapping segments. To remove potential tRNA and rRNA promoter regions or promoter regions without any activity the resulting data frame was merged with a data frame of a previously conducted transcriptome analysis of an RNAseq experiment with *Curvibacter* wildtype cells (Further details s. Pietschke *et al*. 2017 [2]). The merging step eliminated all potential promoter sequences from genes that are not part of the transcriptome analysis, and as a result, these genes do not exhibit any expression levels in the standard R2A growth media of *Curvibacter*. In addition, it excluded all genes labeled as tRNA or rRNA since they were not present in the transcriptome data frame. This step was vital in eliminating conflicting or redundant information within the analyzed gene sequences. In a next step five sequences have been filtered out for containing restriction binding sites for BsaI, BsmBI and BbsI. Sequences containing these binding sites were excluded. After this 722 candidates remained in the data frame. As the sequence synthesis order was capped to 500 sequences this selection was cut further: the 350 smallest sequences included all sequences from 60 to 98 bp were included first, as the sequence synthesis order was capped to a maximum length of 150 bp including added restriction sites on each site. The remaining sequences were sorted by their read counts derived from the RNAseq experiment mentioned above. As the read counts encompassed three orders of magnitudes of read count levels, the 50 5^*′*^ upstream regions from genes with reads of each order of magnitude were picked to cover a wide range of potential expression levels. All sequences were subsequently trimmed from the 5^*′*^ end to reach a length of 98 base pairs. Restriction sites for golden gate cloning were then added, followed by the insertion of random bases behind the restriction site to meet the synthesis specifications, ensuring that all sequences reached a uniform length of exactly 150 base pairs.

### 2.2 Transcriptome Read Mapping

Raw RNAseq reads were obtained as described in Pietschke *et al*. 2017 [2]. Before mapping the obtained sequences against the reference genome of *Curvibacter*, the sequences were subjected to quality control and preprocessing. Sequences were trimmed using trimmomatic (Version 0.39, [18]). Trimmed FASTQ files were analyzed for quality using FASTQC (Version 0.11.9, [19]). Subsequently, the preprocessed sequences were mapped against the reference genome of *Curvibacter* utilizing the kallisto mapper (Version 0.50.0, [20]). RAW sequences are uploaded to NCBI as BioProject PRJNA1082616. They will be made public after full publication and are available as of now upon request. The relevant code can be found in this GitHub repository: (https://github.com/Kanomble/curvibacter_transcriptomics).

### 2.3 Library Golden Gate Cloning

40 ng/kb of DNA library (but a minimum of 5 ng per reaction) were added to 20ng/kb of entry vector to maximize yield of successful integration without compromising efficiency. Golden Gate Reaction was performed according to standard protocol, but the final digest duration was increased to 1 h to reduce entry vector religation. 50 μl of highly competent Dh5 Alpha cells were transformed with 10 μl of Golden Gate reaction and plated on 4 selection plates to reduce colony crowding. After 24 h of growth, a minimum of 5.000 (for a library with 500 sequences) colonies of equal size were obtained and scraped off the plates. Plasmid DNA was extracted from the cell mixture using a standard MiniPrep kit from Macherey Nagel. The *E. coli* donor strain for conjugation into *Curvibacter* was retransformed with the plasmid library to yield a minimum of 5.000 colonies.

### 2.4 Conjugation of Library Vectors into *Curvibacter* sp. AEP1-3 glmS::GFP

*Curvibacter* with a GFP insertion in the glmS locus was inoculated from a fresh plate into R2A+ media and grown for 36h to stationary phase prior to conjugation. DAP auxotroph *E. coli* donor cells were directly scraped off transformation plates, washed in LB media and used for conjugation.

3 ml of *E. coli* donor cells at OD1 and 5 ml of *Curvibacter* stationary phase at OD2 were mixed and the conjugation mix was centrifuged at 5000 rpm for 5 min. Cells were washed in 1 ml of R2A+ and centrifuged as before. Cells were resuspended in 100 μl of R2A+ and spotted on 4 plates of R2A media without the addition of DAP or antibiotics.

Plates were incubated overnight at 30°C and cell spots were scraped off and washed in 1 ml of R2A+. 50 μl of the conjugation mixture was separately plated on a R2A plate containing the respective antibiotic for quality control. The rest of the mixture was centrifuged, resuspended in 200 μl of the remaining media and spread equally over 8 R2A plates containing the respective antibiotic. If the colony count on the quality control plate was higher than 250, the total conjugation yielded over 5000 conjugation events and the plates could be used further. Conjugation plates were scraped and *Curvibacter* cells were washed in 1 ml of R2A+. Cells were diluted to OD 0.02 in R2A and sorted as described below.

### 2.5 Flow cytometry and Cell sorting

*Curvibacter* cells were sorted using the CytoFlex SRT Benchtop Cell Sorter. Forward scatter (FSC) and side scatter (SSC) was measured using a 488 nm laser and a 488/8 nm Bandpass filter. Violet side scatter (VSSC) was measured using a 405 nm laser and a 405/5 nm Bandpass filter. GFP fluorescence was measured using a 488 nm laser and a 525/40 nm Bandpass filter. mCherry fluorescence was measured using a 561 nm laser and a 610/20 nm Bandpass filter. Following gain settings were used:

**Table 1:** Gain settings for flow cytometry.

| Filter | Gain setting (X/3000) |
| --- | --- |
| FSC | 76 |
| SSC | 299 |
| VSSC | 106 |
| GFP | 196 |
| mCherry | 1216 |

The cell population of interest was sorted based on a mCherry fluorescence higher than the background signal. To determine the gate for background fluorescence, the background strain *Curvibacter* AEP1-3 glmS::GFP was used and the gate was set to exclude 99% of this population. The subpopulation was split into four quadrants based on their GFP and mCherry signal: green fluorescence from the genomic GFP over 100.000 AU was considered “high green” (HG), below was considered “low green” (LG) fluorescence. Equally, red fluorescence above 7.000 AU was considered “high red” (HR), below was considered “low red” fluorescence. The quadrant of each sorted cell is stated in the inventory list (s. Supplementary Table 2). 2000 Cells were sorted into 4 tubes containing 100 μl of R2A based on their combined red and green fluorescence signal.

This mixture was plated on R2A plates and incubated at 30°C for 48 h. Colonies of surviving cells were tested for successful promoter integration in the expression vector using cPCR and Sanger Sequencing. Colonies were inoculated in 1 ml of R2A+ containing the respective antibiotic and grown for 48 h. 1 ml of 50% (v/v) glycerol in distilled water was added and cultures were frozen at -80°C for further use.

### 2.6 Bulk Fluorescence Intensity measurements

*Curvibacter* cells containing one of the selected 5^*′*^ upstream non-coding regions were inoculated from a fresh R2A+ plate into R2A+ media and grown for 36 h at 30°C in 24 well plates in a BMG labtech Clariostar plate reader until stationary phase. From these pre-cultures, main cultures were inoculated to an OD of 0.05. Growth and fluorescence was monitored over 36 h. mCherry fluorescence from the reporter constructed was monitored at 570/15 nm bandwidth excitation and 620/20 nm bandwidth emission and GFP fluorescence from the genomically integrated GFP was monitored at 470/15 nm bandwidth excitation and 515/20 nm bandwidth emission.

### 2.7 *Curvibacter* Growth Media

R2A+ media was prepared by adding additional nutrients to premixed R2A from Carl Roth.

**Table 2:**
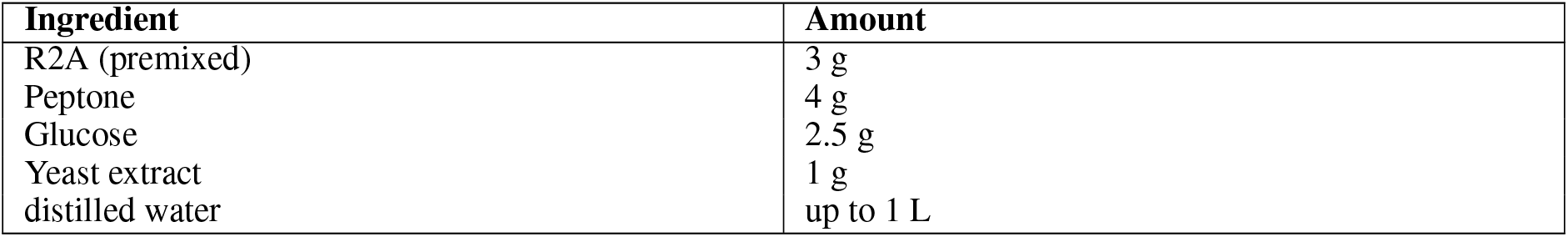
R2A+ media composition.

### 2.8 Mathematical Operations for RFU Assessment

Fluorescence intensity measurements were adjusted to eliminate background signals by employing a media-only control well. The corrected fluorescence intensity for a specific fluorophore was obtained by subtracting the signal in the presence of media control from the signal of that fluorophore alone.

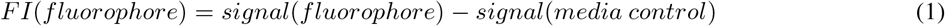

The fluorescence intensity FI(fluorophore) is the value for the Relative Fluorescence Unit (RFU) of the measured fluorophore (mCherry or GFP, signal(fluorophore)) substracted by the RFU of the media control (R2A+, signal(media control).

The datasets obtained from the plate reader (BMG labtech clariostar), which included Biomass and FI data points for mCherry (RFP) and GFP, underwent a filtering process to remove outliers and reduce noise. This smoothing step utilized the savgol_filter function from the Python scipy package, implementing the Savitzky-Golay smoothing technique [21]. The Savitzky-Golay smoothing filter is a data processing method commonly employed in signal processing and data analysis. Its purpose is to smooth noisy data while preserving essential signal features. This step was taken to enhance the accuracy of determinations regarding stationary and exponential growth phases.

The time points for defining the exponential and stationary phases were determined by applying the Savitzky-Golay smoothing technique to the raw OD600 values and calculating the maximum slope for the exponential phase, as well as the maximum OD600 value for the stationary phase. To account for variations in biomass, relative fluorescence units (RFU) were further normalized using the GFP intensity as a reliable proxy for biomass. This normalization was carried out as follows:

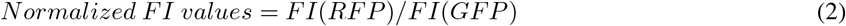

In order to assess changes in promoter activity across different growth phases, FI values were compared between the exponential and stationary phases. This calculation was performed using the formula:

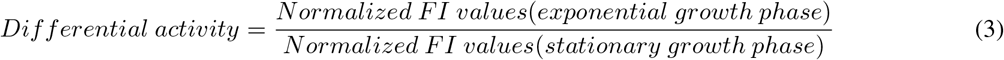

Values below 0.7 indicated promoters more active during the stationary phase, values above 1.3 suggested greater activity in the exponential phase, and values falling in between were indicative of promoters that did not show a significant preference for either growth phase. This method allowed for a comprehensive assessment of promoter behavior in relation to different growth phases.

### 2.9 Inference of Transcription Factor Binding Motifs

To identify potential transcription factor binding sites and motifs (TF-Motifs), the nucleotide sequences of the 33 5^*′*^ upstream non-coding regions identified via Flow Cell Cytometry were used as input sequences for the XSTREME algorithm ([22]). The XSTREME algorithm conducts a comprehensive motif analysis. It was configured with default settings to search for binding motifs within the prokaryotic CollectTF database (http://www.collectf.org/browse/home/), which contains known bacterial transcription factor binding sites.

## 3 Results

### 3.1 Development of a streamlined workflow for promoter mining

We designed a synthetic biology approach for promoter mining in sequenced bacterial species. The details of the workflow to filter potential promoter sites are described in the first methods section in detail. Briefly, candidate promoter sequences for this study were directly harvested from the *Curvibacter* genome sequence. As the *Curvibacter* genome contains 4096 predicted genes, nearly the same amount of intergenic regions (as some genes overlap) exist as potential promoter sites. Filtering steps removed intergenic regions of divergent genes; with a length <60 and >170bp; promoter sites of tRNA and rRNA genes as well as those intergenic regions containing restriction sites of BsaI, BsmBI and BbsI. A table of all ordered sequences is available in Supplementary Table 1. The complete workflow is available on Github (https://github.com/Kanomble/curvibacter_promotor_studies) and can be adapted to any bacterial genomic sequence. In the following we show how representative our selection is for the genome of *Curvibacter* sp. AEP1-3.

### 3.2 Representativeness of extracted 5′ upstream non-coding regions in *Curvibacter* sp. AEP1-3

The 33 5^*′*^ upstream non-coding regions that were confirmed by Flow Cytometry (see next section) are evenly distributed throughout the entire *Curvibacter* genome (s. Figure 2). The length of the confirmed 5^*′*^ upstream non-coding regions ranges from 60 bp to 146 bp. Further, sequences of these confirmed candidate sequences were analyzed for the occurence of common motifs (s. Figure 3).

Among the 33 5^*′*^ upstream non-coding regions, five distinct transcription factor binding site motifs (TF-Motifs) have been identified using the XSTREME algorithm from the MEME-suite portal, with a setup that enables searching for TF-Motifs within the CollectTF database for bacterial transcription factor binding sites [22]. The identified motifs are found in several transcriptional regulators. For instance, the first motif (1) exhibits similarities to TF-Motifs of the AmrZ and LasR genes of *Pseudomonas aeruginosa* [23, 24]. AmrZ serves as a transcriptional activator and/or repressor of virulence factors, as well as genes involved in environmental adaptation. LasR, on the other hand, serves as a transcriptional activator of the elastase structural gene LasB [25] and it is considered as a transcriptional activator for virulence genes in *Pseudomonas aeruginosa* [26]. In addition, LasR is a homolog to the CurR1 and CurR2 genes in *Curvibacter*, which are described in Pietschke *et al*. [2]. In 23 of 25 5^*′*^ upstream non-coding region strains that were further analyzed by bulk fluorescence measurement (further referred to as candidate promoters) (s. Figure 4a) at least one of the inferred TF-Motifs has been identified (s. Supplementary Table 3). A detailed description of all identified motifs can be found in Supplementary Table 3.

### 3.3 *Curvibacter* strains carrying functional reporter constructs show a range of expression levels

33 unique candidate promoters sorted by flow cytometry were further analyzed for their expression level throughout different growth phases (s. Figure 4a) using bulk fluorescence measurement in a plate reader (full list of candidates s. Supplementary Table 1). The candidates vary in length and GC content. For instance, CPL0025 has a length of 76 bp with a GC content of 40%. In contrast, CPL0095 is 113 bp long with a GC content of 42%, and CPL0022 spans 123 bp with a GC content of 46%. From 33 total sorted candidates, 25 showed detectable expression levels and were therefore included in the following analysis (further referred to as candidate promoters).

All strains are based on the same *Curvibacter* background strain containing a genomic GFP integration in the *glmS* locus with a constant expression level relative to biomass until early stationary phase (s. Supplementary Figure 2). As this GFP signal was less noisy compared to OD measurements at optical densities near OD 0.1 and as the fluorophore accumulation in the late stationary phase due to protein aggregation was nearly identical for both the reporter mCherry construct and the genomic GFP integration, the GFP signal was further used as a normalization factor for the RFP signal (s. equation three in Mathematical Operations for RFU Assessment). This reduces noise in low optical density cultures and leads to a stable signal in the later stationary phase, making it easier to determine reporter activity during exponential- and stationary phase. Relative fluorescence units are therefore given as the fraction of RFP/GFP intensity. Supplementary Figure 2 shows the correlation between GFP and biomass of the background strain, showing that GFP expression remains constant relative to biomass until stationary phase is reached and linearly increases after reaching stationary phase in the same way that it does in in trans expression systems, effectively negating this drift. Figure 5 a-d shows a linear relation for RFP/GFP during the stationary phase as a result of this.

Figure 4a shows the relative fluorescence units (RFUs) of mCherry normalized to GFP for all candidates, which showed expression levels above the background noise. The above values serve as a proxy for the relative expression levels of their corresponding promoter sequences during the exponential phase and the stationary phase and should guide investigators in picking expression systems for their specific use case. Expression levels range from 0.61 to 0.005 relative to GFP in the stationary phase, encompassing two orders of magnitude in terms of expression strength. Among the 25 analyzed candidate sequences, 12 show less than 75% of activity during the exponential phase compared to the stationary phase, while 10 display relatively consistent expression strength regardless of the current growth phase (Figure 4b). Additionally, three candidate sequences demonstrate at least 1.25-fold higher activity during the exponential growth phase than in the stationary phase.

**Figure 2:**
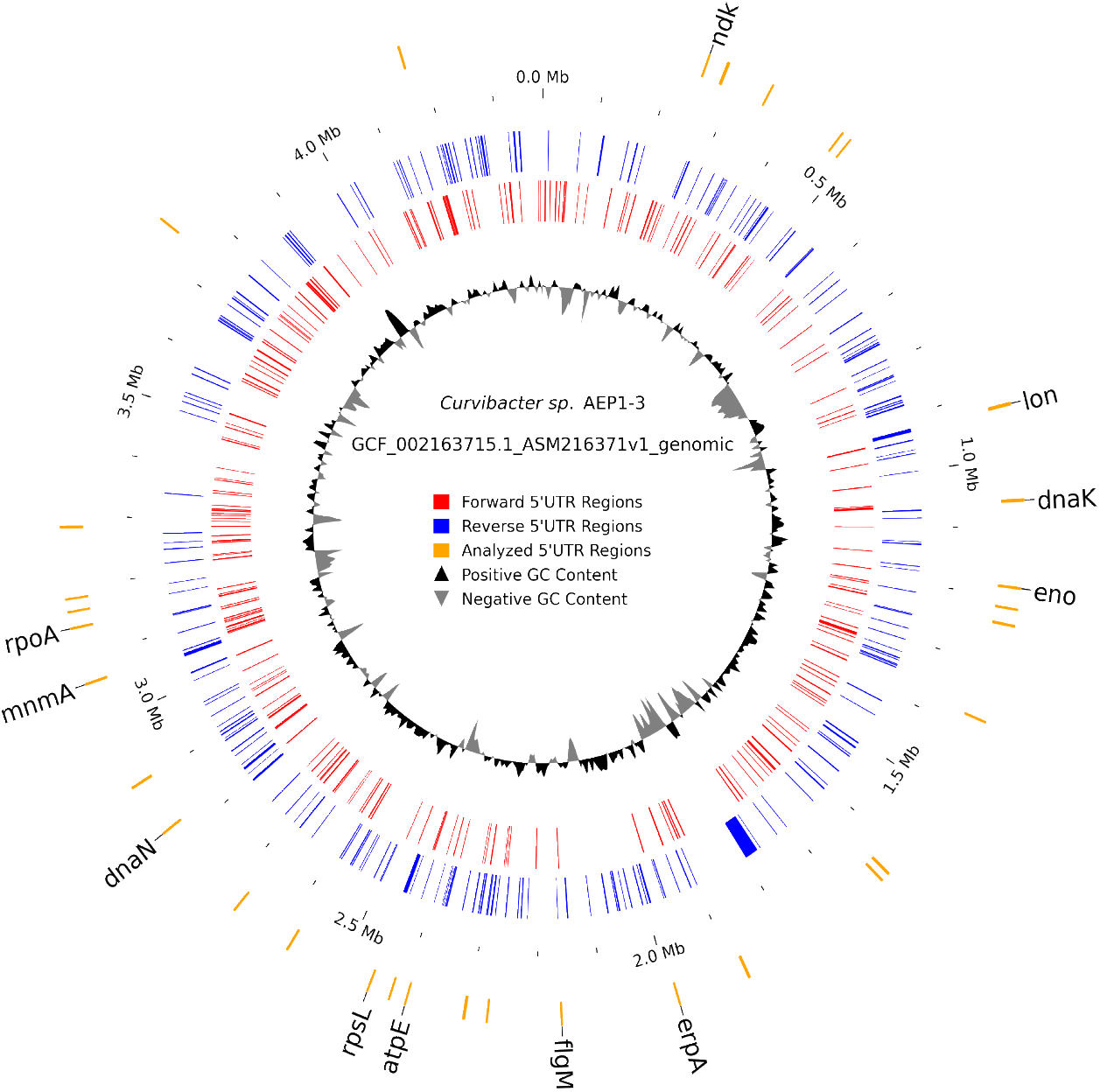
Distribution of extracted 5′ upstream non-coding regions within the genome of *Curvibacter*. The innermost circle is composed of a density plot that showcases the GC content of the respective genome regions. The two following red and blue circles highlight the initially extracted 500 5^*′*^ upstream non-coding regions. Blue lines correspond to genes with a forward orientation (clockwise), red lines vice versa. The outer circle represents the 33 via Flow Cytometry sorted 5^*′*^ upstream non-coding regions. 5^*′*^ upstream non-coding regions of CDS regions labeled as “hypothetical protein” or with protein names that are too long are not labeled.

**Figure 3:**
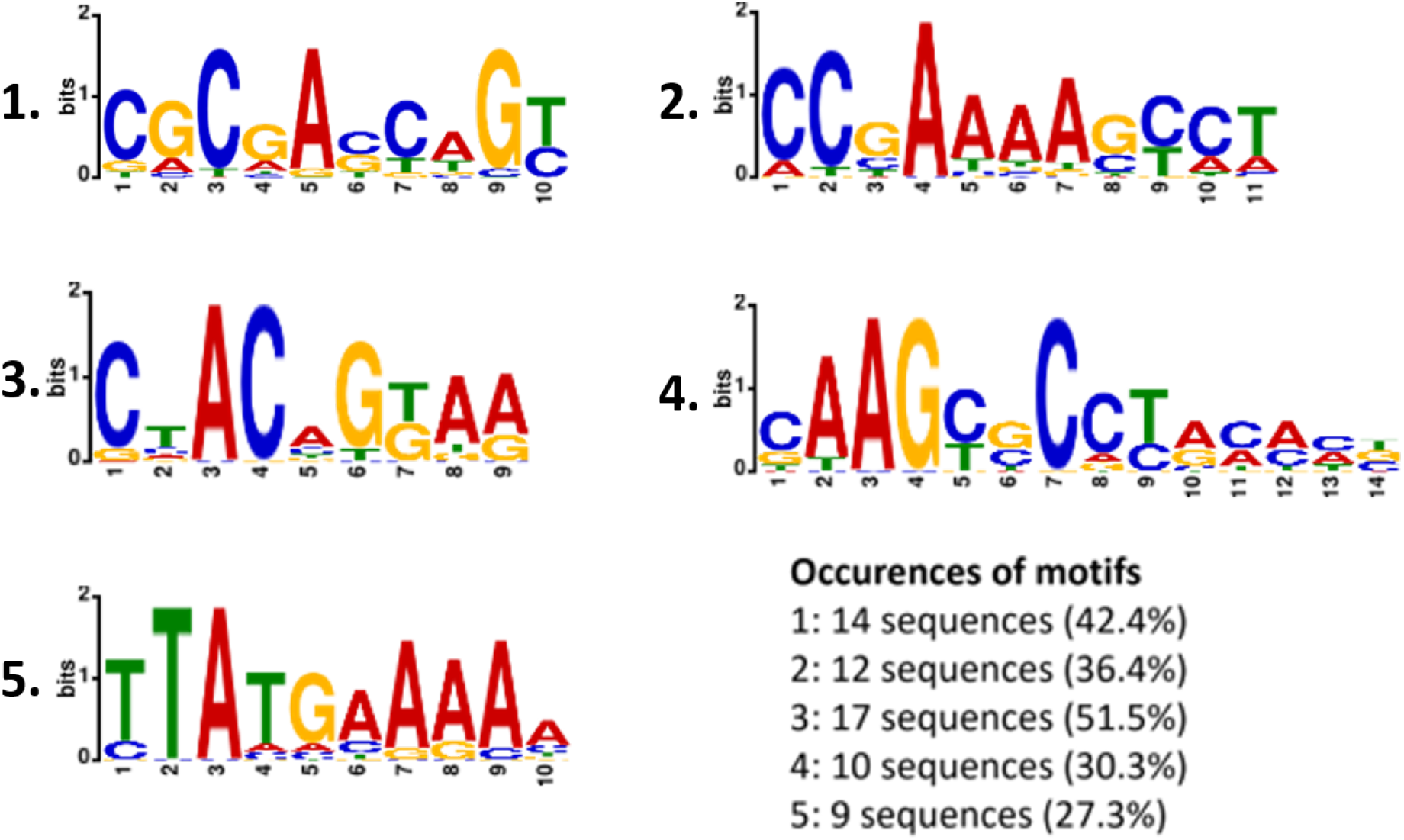
Conserved sequence motifs within the analyzed 33 5′ upstream non-coding regions of *Curvibacter*. Sequence motifs are part of the CollectTF database. Figure highlights the 5 most prevalent motifs, additional motifs can be found in Supplementary Table 3.

**Figure 4:**
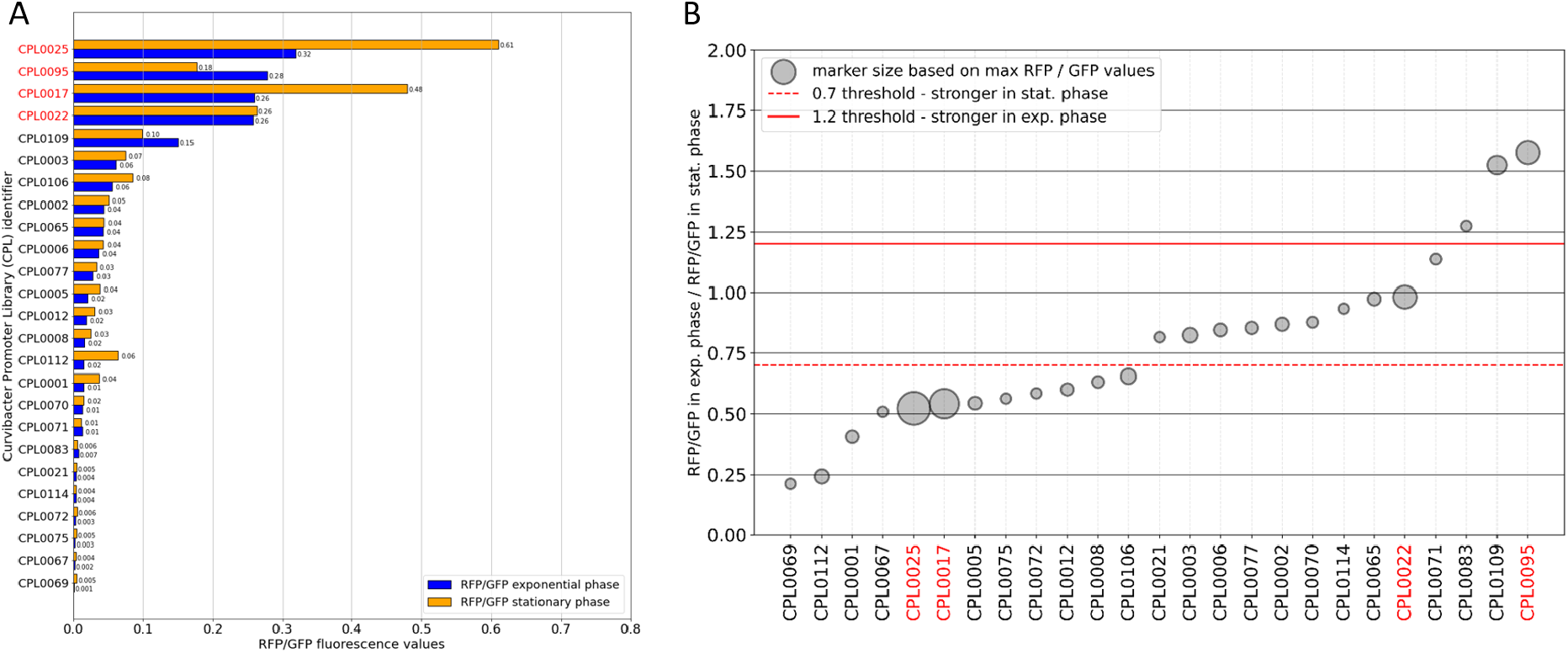
Left: Promoter expression levels as measured by RFP/GFP levels within the exponential growth phase and stationary growth phase. Promoter sequences are sorted in decreasing order based on the RFP/GFP value during the exponential growth phase. The red marked CPL identifiers represent the highlighted sequences in the section below. **Right: Exponential phase and stationary phase proportion of promoter expression strengths**. Promoters with values under 0.7 are more active during the stationary growth phase, while promoter sequences with values above 1.2 are considered to be more active during the exponential growth phase. Dot size correlates with overall expression level. The red marked CPL identifiers represent the highlighted sequences in the section below.

### 3.4 Activity level of candidate promoters shows distinct temporal expression dynamics over growth phases

In this section we provide detailed information of the measured fluorescence activity over time of three (plus CPL0017 as control) selected candidate promoters of *Curvibacter*. We recommend these promoters for further experimental use as they cover a range of different temporal expression patterns and strengths.

The CPL0025 (Figure 5B) sequence is the promoter of the gene AEP_RS11205, annotated as the acyl-homoserine-lactone (AHL) synthase (RefSeq protein identifier: WP_232459811) and described as Curl2 in Pietschke *et al*. [2]. The full promoter region (519 bp) of the AHL synthase Curl2 is activated by homoserine lactones, a bacterial quorum sensing molecules which plays a crucial role in regulating gene expression in response to population density [27]. Here we show that even the smaller promoter region of 76 bp could drive the expression of our reporter construct. The 5’UTR shows lower expression levels during exponential growth and elevated expression levels after entering stationary phase. CPL0025 was the strongest candidate among all tested promoters, surpassing even the expression level of the highly active J23100-RBS* promoter (CPL0017), which was used as reference (Figure 5a). A TF-Motif similar to the AHL activated transcriptional regulator LasR (s. Supplementary Table 3) was not detected within the 76 bp long 5’UTR of CPL0025. The only motif found in this 5’UTR is similar to TF-Motifs found in Gram-positive bacteria, the motif is located within the positions 20 - 29 (TTACAAGAAA) of the 5’UTR. Specifically, similar to motifs of the global transcriptional regulator CodY from *Lactococcus lactis* and *Streptococcus pyogenes* as well as for CcpA from *Streptococcus pneumoniae* (s. Figure 3 5. and Supplementary Table 3) [28, 29, 30, 31].

**Figure 5:**
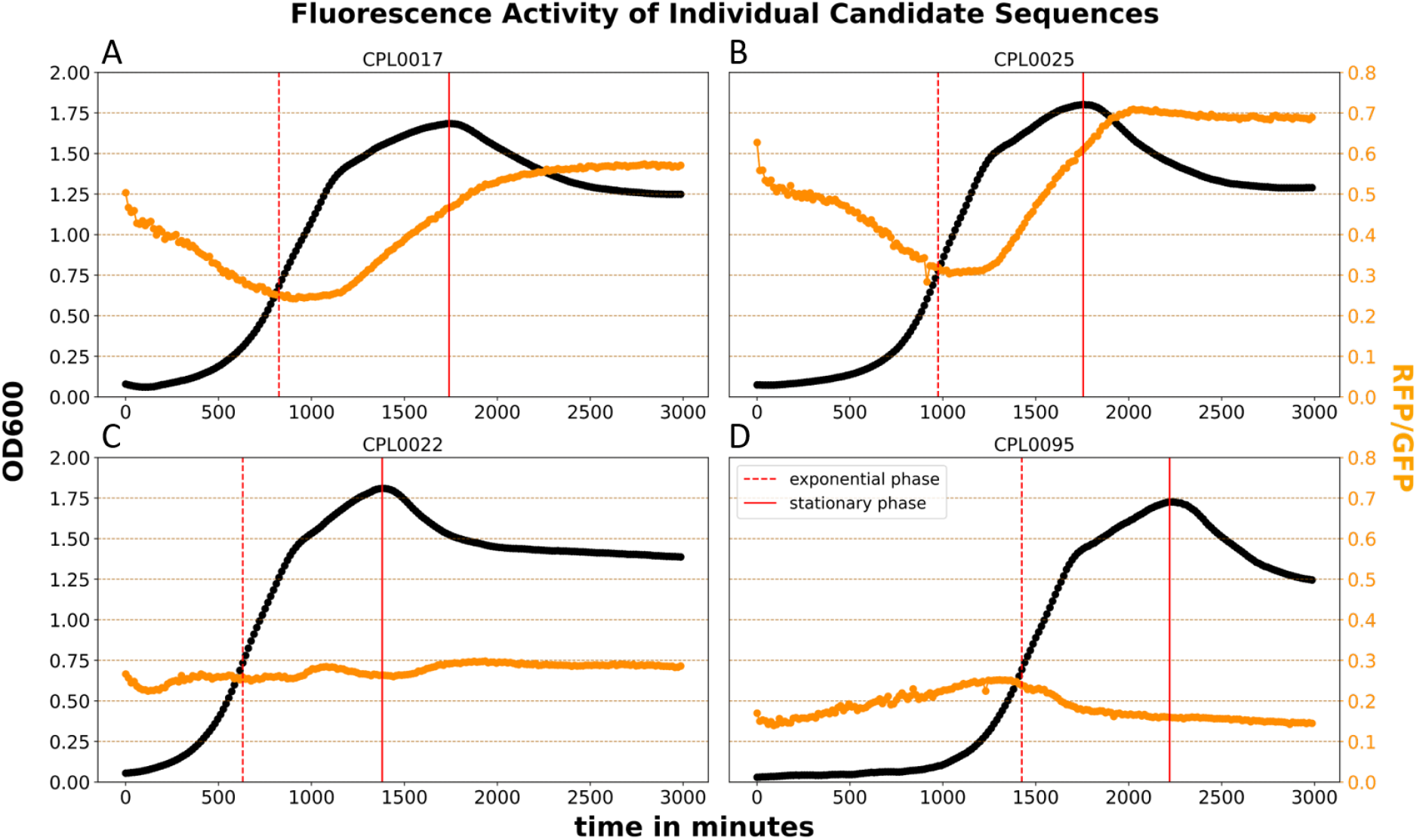
Promoter expression levels for the highlighted sequences CPL0017 (control), CPL0025, CPL0022 and CPL0095. Full and dotted lines represent time point of exponential and stationary phase, respectively, which was used for Figure 5B. RFU (orange line) equals RFP intensity (candidate promoter) normalized to GFP intensity (which is located on the genome).

Sequence CPL0022 (Figure 5c) is the promoter of the gene AEP_RS05045 expressing DnaK (RefSeq protein identifier: WP_087494375), a molecular chaperone protein of the (Heat-shock-protein 70) Hsp70 family. The *dnaK* candidate promoter shows a very constant expression level throughout all growth phases in *Curvibacter* compared to all other tested promoters, with minor bursts of transcriptional activity during late exponential and early stationary growth phases. In comparison with other sequences in this study, the *dnaK* shows a very constant expression level throughout all growth phases shows a relatively high expression level and very little bias towards growth phases. CPL0022 contains 9 bp long TF-Motif for LasR binding within the positions 12 - 21 (CACAACCAGC) of the 5’UTR sequence. Additionally, CPL0022 contains a CodY motif from *Bacillus anthracis* and a ExpR motif from *S. melliloti* (s. Supplementary Table 3).

CPL0095 (Figure 5d) is the promoter of the gene AEP_RS11420 (RefSeq protein identifier: WP_011466063) expressing RpsL, a 12S protein component of the 30S ribosomal subunit. This candidate promoter displays high activity during the exponential phase, with a steady increase in activity until the mid-exponential phase. The activity then decreases to approximately half of its maximum during the stationary phase. CPL0095 contains a range of sequence motifs, such as a LasR motif from *P. aeruginosa*, CodY from *B. anthracis* as well as *S. pyogenes*, and a LexA motif from *V. parahaemolyticus* (s. Supplementary Table 3).

### 3.5 Comparison of RNAseq- and RFU-assessed promoter strength only shows a weak correlation

Figure 6 presents a comparison between the measured and normalized RFU values of the promoter sequences and their corresponding mean read counts across *Curvibacter* RNA-seq samples. When comparing our data on the promoter strength of the candidate sequences to the RNA-seq read counts of the same genes, significant disparities in relative strength within these datasets become evident. The Pearson correlation analysis resulted in an R-value of 0.13, indicating only a weak positive correlation between both datasets. Data points close to the regression line represent sequences that show expression levels in our assays matching closely to the read counts gathered from the RNA-seq analysis, relatively to all data points. For data points falling below the separation line, promoters were stronger in the experimental assessment with respect to their mean read counts in the RNA-seq, and vice versa for data points above the regression line.

**Figure 6:**
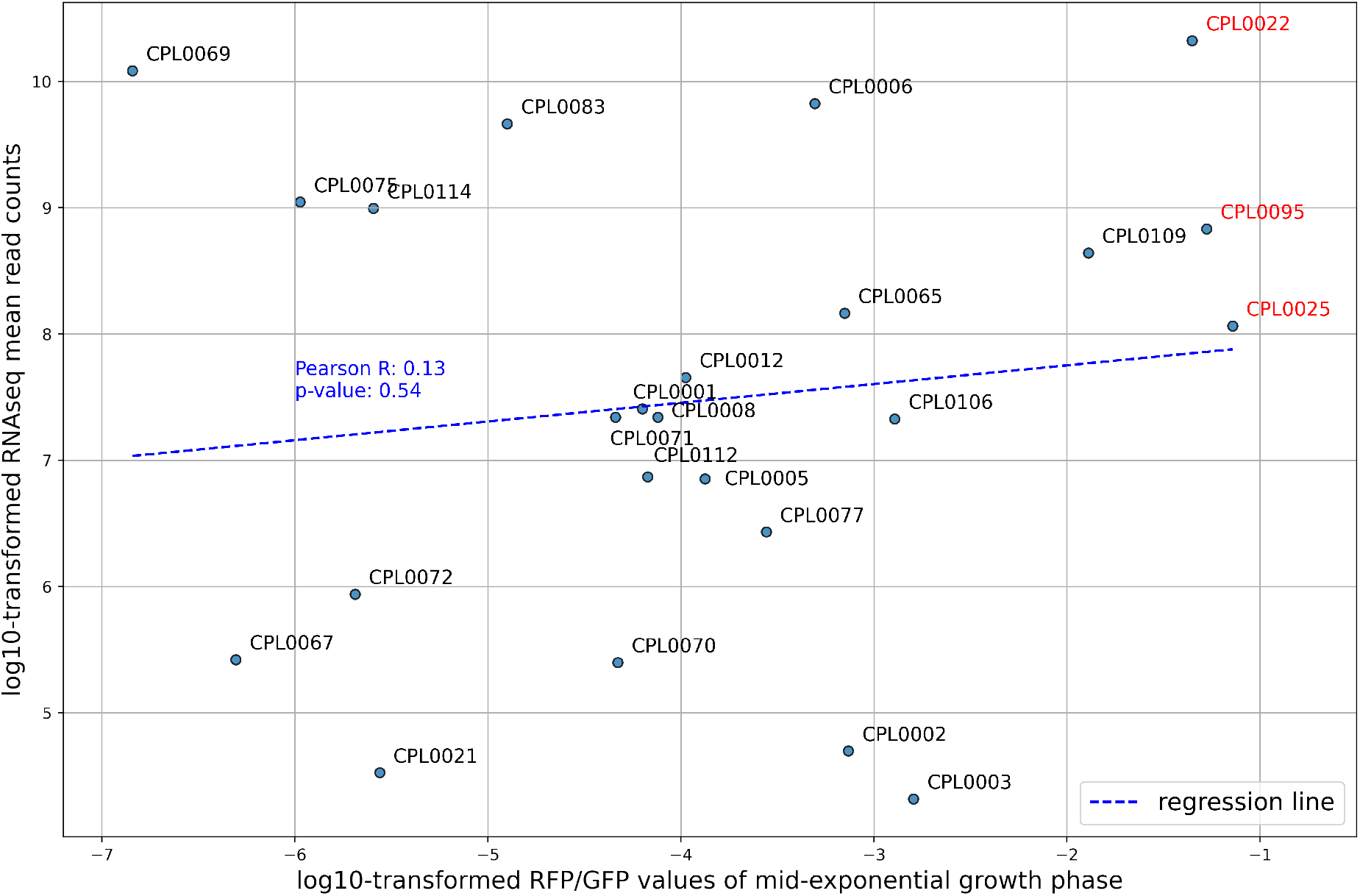
Comparison of promoter strengths in the exponential phase with their corresponding mean RNAseq read counts for candidate promoters. The dotted blue line represents linear regression across all data points. Red marked CPL identifiers represent sequences highlighted in Figure 5.

## 4 Discussion

### 4.1 25 novel promoters for the use in *Curvibacter* show distinct temporal expression dynamics

As *Curvibacter* is a promising model organism we set out in project this project to extract novel expression systems for this species from a self designed oligonucleotide library. This library was generated by mining the *Curvibacter* genome for potential promoter sites in an automated fashion. Positive candidates were first picked via Flow Cytometry and subsequently individual sequences were analyzed by bulk fluorescence measurement. From our 500 initial candidate sequences we found 25 positive candidates that showed expression based on our reporter plasmid. Among these, we could find expression levels over two orders of magnitude and a variety of different temporal expression dynamics over growth phases (s. Figure 4a). We found 12 candidate promoters which show a higher expression level in the stationary phase compared to the exponential phase. 10 candidate promoters showed very little discrimination between growth phases, maintaining a stable level of expression throughout the observed duration and three candidate promoters showed a higher expression level in the exponential phase compared to the stationary phase (s. Figure 4b).

Not only can these new expression platforms be used as tools for expression during different growth phases in liquid medium, but the expression strength assay may also indicate the temporal expression dynamics of these genes in their native genomic context: As expected, many of the candidate promoters with higher activity levels in the exponential phase belong to genes expressing proteins involved in central metabolism and proliferation (50S ribosomal protein L25/general stress protein Ctc, RpsL, Ndk (s. Figure 4b)).

This is in accordance with previous findings which show that bacterial cells are able to recall distinct global expression patterns based on their stage of growth by the spatio-temporal regulation of chromosomal macrodomains [32]. While replication induced transient changes in actual copy numbers are a factor directing genomic transcription biases along the oriC/ter axis [33], the regulation of macrodomains occurs for functionally similar genes through direct DNA topology and transcriptional control. While *in trans* expression systems are per definition not affected by positional effects of the promoter of interest (as they are taken out of their natural, genomic context), they are partially affected by DNA topology [34] (e.g. plasmid supercoiling) and fully affected by transcriptional modulation, under the condition that the entire sequence relevant for regulation is included in the expression system. On the other hand, we see a variety of (often hypothetical proteins) gene functions associated with the candidate promoters where the expression levels are higher in the stationary compared to the exponential phase. This is a result of the general expression bias in plasmid-based expression systems, which tend to exhibit higher expression levels in the stationary phase. Consequently, a bias towards stationary phase expression can be observed, complicating the interpretation of the native context of these genes and their temporal expression dynamics. This effect is primarily attributed to the enrichment of plasmid copy numbers in the stationary phase relative to the number of cells [10, 11, 12]. While saturation of protein density was normalized in our assay by utilizing GFP FI values as a normalization factor for mCherry FI values, a bias introduced due to plasmid copy number enrichment is not. We showed that by harvesting 5^*′*^ upstream non-coding regions from the target species genome we were able to create expression systems that behave differently from most synthetic, orthogonal *in trans* expression systems. These expression systems can now be used to further study *Curvibacter* sp. AEP1-3.

The intial library encompassed 500 5^*′*^UTR sequences from the *Curvibacter* genome. As 25 of these showed detectable expression levels, the discovery rate is therefore at a minimum of five percent (5%). Many potential promoters may not be active under the artificial laboratory environment and hence show little activity, especially considering that R2A is a complex media that already serves a lot of metabolites and thus requires less *de novo* synthesis of many compounds. To eventually raise the success rate of promoter prediction before manually curating the oligonucleotide library, stretches of sequences around the extracted loci could be used as input sequences for a neural network trained by known promoter sequences such as sequences from the PDD [16]. A similar approach was recently conducted by Seo *et al*. for the cyanobacterial species *Synechocystis* sp. PCC 6803 [35]. The AI generated prediction could further be used to extract and construct more efficient oligonucleotide sequences. These sequences can be based not only on a continuous DNA-sequence between gene regions but also on specific k-mers of 5^*′*^UTR sites. Thus motifs responsible for RNA-polymerase recruitment can be located upstream of the sequences ranging into the next gene sequence, which our approach currently does not cover.

### 4.2 Temporal expression dynamics of highlighted promoters may correspond with their biological functions

For applications where a stable expression level is essential or accumulation of protein aggregates is a known issue, we recommend the use of the CPL0022 promoter, which drives expression of the *dnaK* gene in *Curvibacter* [36]. DnaK in *E. coli* is constitutively expressed throughout all of its life cycle and the same seems to account for the DnaK equivalent in *Curvibacter* (s. Figure 5c). The DnaK protein is a molecular chaperon, a class of enzyme involved in guiding correct folding after translation as well as for already matured proteins. While this maintenance is required constantly, it is generally upregulated when bacteria face external stresses that lead to rapid protein degradation such as heat shocks. Thus, DnaK in *E. coli* is part of the Hsp70 protein group. In *Curvibacter*, the CPL0022 promoter also showed a relatively stable expression level throughout all growth phases (s. Figure 5c). It would be interesting to see whether this promoter could be utilized as an inducible expression system by applying heat shocks to the cells as a stimulus, effectively acting as an inducible promoter. As *Curvibacter* is studied due to its symbiotic partnership with its host *Hydra vulgaris*, it would be interesting to see whether this promoter also maintains stable activity when growing on the glycocalyx of *Hydra*.

Alternatively, protein aggregation can also be prevented by using a promoter with lower expression levels during the stationary phase such as the CPL0095 promoter. In its native context, this CPL0095 expresses RpsL, a 12S ribosomal protein of the 30S subunit. This 12S subunit is added late in the biogenesis of the 30S subunit and is essential [37]. Due to the high demand for protein expression during the exponential phase, genes involved in protein translation are upregulated during that phase, explaining the unusual temporal expression dynamic of this candidate promoter (s. Figure 5d). While the expression level of CPL0095 during the exponential growth phase is equal to the strongest sequence CPL0025, it is almost five fold weaker during the stationary phase in comparison (compare to Figure 5b). Further modification to reduce or enhance the general expression level of this promoter could fine tune its function for application in continuous cultivation systems such as chemostats. It is likely that other promoters driving gene expression of proteins involved in the assembly of ribosomal subunits or translation are upregulated in a similar fashion and could be a first avenue to find more promoters that behave similarly to CPL0095.

For high levels of expression we recommend the use of CPL0025 (s. Figure 5b) or CPL0017 (s. Figure 5a) which has been used in this study as a reference sequence. CPL0025 had the highest level of expression among all tested sequences. Both promoters are well suited for the expression of proteins with very little burden on the host cell metabolism, such as fluorophore proteins for imaging. The gene expressed from CPL0025 functions as an AHL synthase (Curl2), as described by Pietschke et al. [2]. They have shown that the full promoter region (519 bp) of CPL0025 is activated by AHLs produced by *Curvibacter*, as well as by AHLs modified by *Hydra vulgaris*. Here we show, that the shortened promoter region of 76 bp is able to drive a strong expression of our reporter (s. Figure 5b). Within the identified transcription factor binding motifs, a TF-Motif similar to a motif discovered for the transcriptional regulator LasR of *Pseudomonas aeruginosa* has been identified [23]. LasR is a LuxR-type regulatory protein and a key component in the quorum sensing system of *Pseudomonas aeruginosa*. LasR binds to AHLs activating the expression of genes involved in various virulence factors and genes important for the adaptation to the environment [38]. However, no LasR TF-Motif can be found within the CPL0025 5’UTR but a binding motif for the transcriptional regulator CodY from *Lactococcus lactis* and *Streptococcus pyogenes* as well as for CcpA from *Streptococcus pneumoniae*. Those transcriptional regulators are known to regulate the expression of a wide range of genes, e.g. genes responsible for carbohydrate and (p)ppGpp metabolism or virulence factors [29, 30, 31]. Regarding the expression of the reporter driven by CPL0025 and the previous finding that the promoter region of Curl2 is activated by AHLs, it is possible that the motif responsible for AHL induction is located within the 76 bp long 5’UTR of CPL0025. It is unclear whether the high expression level of CPL0025 in the stationary phase are result of transcriptional changes directly related to the growth phase or a result of increasing levels of AHLs in the media or both.

### 4.3 RNAseq data alone is insufficient to predict the usability of promoters for expression platforms but can aid in selecting best candidates

For the 25 promoters that showed notable activity in our assay (s. Figure 4a), we compared the expression level to the RNAseq mean read counts (s. Figure 6). While some promoters, especially around the midrange of transcriptional strength in the RNAseq data set, align very well with the experimentally verified expression strength, in many cases both data sets do not seem to match very well, indicating that the RNAseq results could only partially predict the expression levels in our assay. While for some sequences the experimentally assessed expression level was higher than expected given the low read counts (eg. sequences CPL0003, CPL0002, CPL0025 in Figure 6) for other sequences the opposite was the case (eg. sequences CPL0069, CPL0075, CPL0114, CPL0083, CPL0006 Figure 6).

As the growth conditions between the RNAseq experiment and the reporter activity assay deviated (18°C growth in R2A for the RNAseq data set and 30°C in R2a+ media), we can assume that in some cases temperature sensitivity of gene activity may play a part in the overall noise between data sets. However, the activity of the *dnaK* promoter is comparably strong in the RNAseq data set (s. Figure 6) which was taken at 18°C, indicating that heat stress induced signals may not be a factor affecting expression levels too much. We can’t exclude transcriptional changes due to heat stress responses or elevated growth rates as a reason for the deviations observed between RNAseq data and our bulk fluorescence measurements. However, other factors may also contribute to the low comparability of the two data sets. It is important to mention that while RNAseq data reports on the relative abundance of cellular mRNA level, it acts as a proxy for transcriptional strength levels of genes. Reporter gene activity however is measured as the amount of active reporter protein; mCherry in our case; and acts therefore as a coupled proxy for transcriptional as well as translational strength levels. Comparison of such data always has to consider the fact that large discrepancies in translational strength levels of different mRNAs due to different ribosomal binding site affinities may have a huge impact on the abundance of gene activity on mRNA and protein level. Lale *et al*. also reported discrepancies between measured mRNA levels and protein levels of promoters from their GeneEE library [15].

We could find some sequences to have almost no reporter activity despite high read counts in our RNAseq data. While we cant exclude truncation of sequence elements vital for transcription in some cases, a weak translation level may be another reason for the low reporter activity. On the other hand, CPL0025 for instance shows a surprisingly high reporter activity despite moderate read counts in the RNAseq data. Even though unlikely, it is entirely possible that a regulatory site may actually be located upstream of the intergenic region, either as part of the proceeding gene or even further, repressing this promoter in its native context, which has not been integrated into the reporter expression system.

Concluding, we can therefore not claim a strong correlation between RNAseq data and performance of the reporter plasmid. To reinform our pipeline based on these results, we can conclude that while picking candidates based on high RNAseq read counts may improve the chance to obtain reporter constructs of high activity, it is definitely not a guarantee for success. It also shows that simply picking a strong promoter from the genome and trying to utilize it in an *in trans* context is not guaranteed to yield a strong expression system for a new model organism, again highlighting how essential broader high throughput strategies such as the one employed in our case are for tool development for lesser researched species.

## 5 Outlook

We created a scalable pipeline for the semi-automated discovery of expression systems that can be applied to any bacterial species of interest with an available genome sequence and expression vector. Our workflow was able to find novel expression systems for *Curvibacter* (s. Figure 5) which can now be utilized in a variety of applications. We also showed, that RNAseq results were insufficient to predict a promoter candidates performance in an *in trans* context. It can be extended to specifically search for inducible promoters, another very relevant manipulation tool for novel model species. First, the entire library could be sorted once under “induced” and once under default conditions and sorted in bulk. After sequencing all promoter regions in the expression vector of both populations, the promoters that appear only in the induced population serve as a list of potential candidates for inducible expression systems. In the same way, this method can be used to search for promoters that are active under any condition of interest e.g., for *Curvibacter* in the presence of the host species *Hydra vulgaris*.

In our curated oligonucleotide library approach, we approximate the rate of RNA polymerase activity by assessing the fluorescence of a reporter protein. In contrast, traditional RNA-seq transcriptome analysis involves counting and comparing mapped read abundances of expressed genes. Nevertheless, both experimental designs share the common goal of detecting the rate of transcriptional activity under specific environmental conditions. Comparing these strategies, we argue that both methods show different scalability towards distinct scenarios: while for some applications, it is desired to find transcriptional changes of a few genes under a multitude of different circumstances, in other cases the focus may be on global changes in transcription levels for only a handful of environmental circumstances. Traditional RNAseq workflows allow for a global analysis of a transcriptome but the amount of samples increases linearly with the amount of observed conditions. To maintain an adequate sequencing depth of each individual sample, this leads to scaling sequencing costs based on the amount of samples. Vice versa, our oligonucleotide based screening workflow can be very well adapted to screen many conditions in a single workflow, as cultivation capacities and the scalability of flow cytometry based sorting are the only rate limiting factors. Another advantage is that once positive candidates are sorted and sequenced, the vectors for further studies are already available.

## 6 Contributions

### 6.1 Author Contributions

Maurice Mager: Conceptualization, Methodology, Investigation, Writing - Original Draft, Writing - Review and Editing, Visualization

Lukas Becker: Conceptualization, Software, Formal Analysis, Investigation, Writing - Original Draft, Writing - Review and Editing, Visualization

Nina Schulten: Methodology, Resources

Sebastian Fraune: Writing - Review and Editing, Funding Acquisition

Ilka Axmann: Supervision, Writing - Review and Editing, Funding Acquisition

## 6.2 Acknowledgements

We thank Timo Minten, Petra Kolkhof and Jay Bathia for their support in this project. The implementation of the cell sorter (CytoFLEX SRT) was kindly supported by Dennis Hasenklever.

