## Supplement for "Oligonucleotide Library Assisted Sequence Mining Reveals Promoter Sequences With Distinct Temporal Expression Dynamics For Applications In *Curvibacter* sp. AEP1-3": Supplement summary.docx

Summary of supplemental files

Supplementary table 1:

list of all 500 promoter sequences as ordered for oligonucleotide library synthesis

Supplementary table 2:

Positive candidate sequences recovered after flow cytometry cell sorting

Supplementary figure 1:

Comparison of change in biomass and GFP expression in Curvibacter AEP1-3 glmS::GFP strain carrying CPL0022 reporter plasmid

Supplementary figure 2:

Comparison of change in biomass and GFP expression in Curvibacter AEP1-3 glmS::GFP background strain

Supplementary file 1:

Genbank file of entry vector for library cloning
