## Supplementary figures and images for "Oligonucleotide Library Assisted Sequence Mining Reveals Promoter Sequences With Distinct Temporal Expression Dynamics For Applications In *Curvibacter* sp. AEP1-3"

### Supplementary figure 1.png

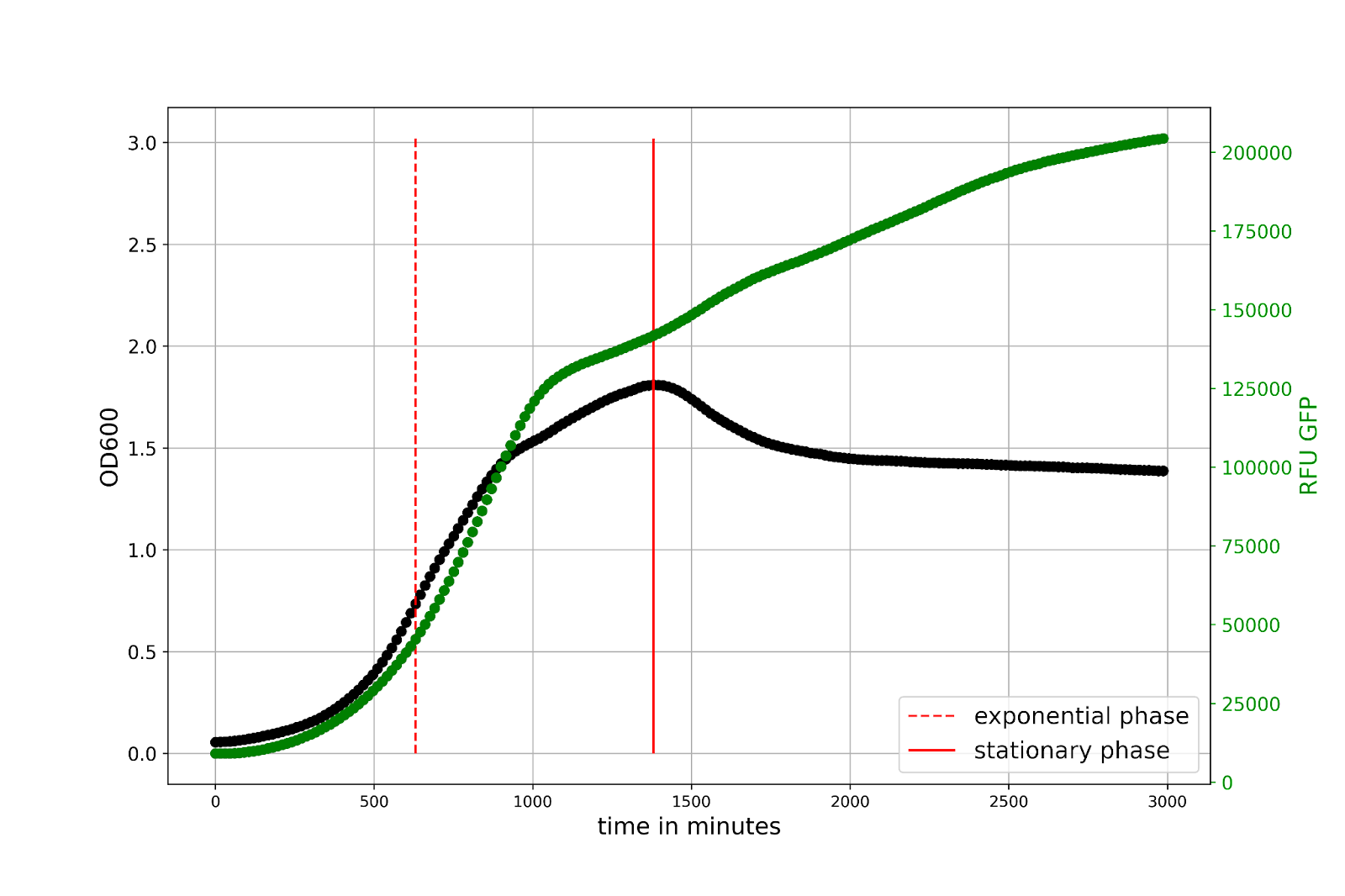

### Supplementary figure 2.png

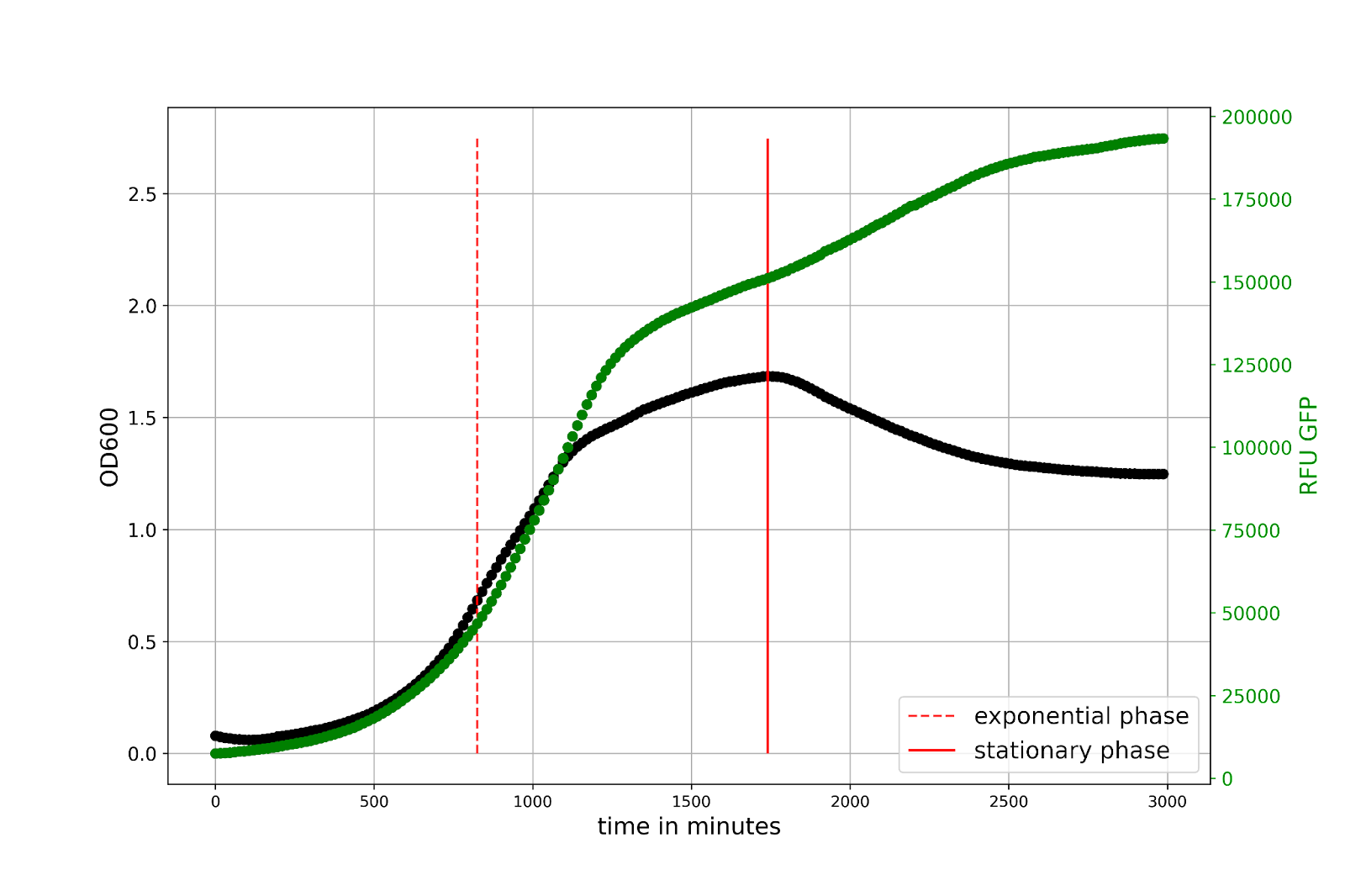
